# Chronic Physical Disturbance Substantially Alters the Response of Biological Soil Crusts to a Wetting Pulse, as Characterized by Metatranscriptomic Sequencing

**DOI:** 10.1101/326181

**Authors:** Blaire Steven, Cheryl R. Kuske

**Affiliations:** Department of Environmental Sciences, Connecticut Agricultural Experiment Station, New Haven, CT, USA.; Bioscience Division, Los Alamos National Laboratory, Los Alamos, NM, USA.

## Abstract

Biological soil crusts (biocrusts) are microbial communities that are a feature of arid surface soils worldwide. In drylands where precipitation is pulsed and ephemeral, the ability of biocrust microbiota to rapidly initiate metabolic activity is critical to their survival. Community gene expression was compared after a short duration (1 hour) wetting pulse in both intact and soils disturbed by chronic foot trampling.

Across the metatranscriptomes the majority of transcripts were cyanobacterial in origin, suggesting that cyanobacteria accounted for the bulk of the transcriptionally active cells. Chronic trampling substantially altered the functional profile of the metatranscriptomes, specifically resulting in a significant decrease in transcripts for nitrogen fixation. Soil depth (biocrust and below crust) was a relatively small factor in differentiating the metatranscriptomes, suggesting that the metabolically active bacteria were similar between shallow soil horizons. The dry samples were consistently enriched for hydrogenase genes, indicating that molecular hydrogen may serve as an energy source for the desiccated soil communities. The water pulse was associated with a restructuring of the metatranscriptome, particularly for the biocrusts. Biocrusts increased transcripts for photosynthesis and carbon fixation, suggesting a rapid resuscitation upon wetting. In contrast, the trampled surface soils showed a much smaller response to wetting, indicating that trampling altered the metabolic response of the community. Finally, several biogeochemical cycling genes in carbon and nitrogen cycling were assessed for their change in abundance due to wetting in the biocrusts. Different transcripts encoding the same gene product did not show a consensus response, with some more abundant in dry or wet biocrusts, highlighting the challenges in relating transcript abundance to biogeochemical cycling rates.

These observations demonstrate that metatranscriptome sequencing was able to distinguish alterations in the function of arid soil microbial communities at two varying temporal scales, a long-term ecosystems disturbance through foot trampling, and a short term wetting pulse. Thus, community metatranscriptomes have the potential to inform studies on the response and resilience of biocrusts to various environmental perturbations.

## Introduction

Drylands are defined by their lack of a vital resource, precipitation. Thus, water availability is a significant force in shaping the structure and function of drylands or arid ecosystems. Biological soil crusts (biocrusts) are a community of cyanobacteria, associated heterotrophic bacteria, fungi and mosses that colonize arid soils worldwide (Belnap, 2003; Belnap et al., 2016; Pointing and Belnap, 2012). These communities inhabit plant interspaces and are important players in fixing carbon and nitrogen in dryland soils (Barger et al., 2016; Evans and Lange, 2001). Because of the unpredictable periodicity and amount of precipitation, biocrusts respond quickly to wetting events to exploit the available moisture to repair cell damage and synthesize new biomass. For example, when precipitation wets surface soils, biocrust cyanobacteria display a property of “hydrotaxis” and move toward the available water (Garcia-Pichel and Pringault, 2001; Pringault and Garcia-Pichel, 2004). Pigments, such as chlorophyll *a,* are rapidly produced, resulting in a “greening” of the surface soils (Abed et al., 2014). Concurrently, consumption of oxygen and evolution of CO_2_ by the biocrusts can be detected, indicating vigorous metabolic activity (Rajeev et al., 2013; Tucker et al., 2017). Wetting pulses can also stimulate “blooms” of specific bacteria, highlighting that specific taxa have differing thresholds and responses to wetting (Karaoz et al., 2018). Despite the multitude of metabolic and physiological shifts associated with wetting in biocrusts, the specific genes and biochemistry underlying these processes are largely unknown.

*Microcoleus vaginatus*, the keystone species of biocrusts, shows significant alterations in gene expression over wetting cycles, as assessed by RNA microarrays. These experiments show that *M. vaginatus* rapidly resuscitates after the addition of water with an upregulation of genes for photosynthesis, anabolic metabolism, and DNA repair (Rajeev et al., 2013). Similarly, the desert moss *Syntrichia caninervis* increases gene expression for photosynthesis, membrane integrity and protein synthesis upon wetting (Gao et al., 2014). However, these results consider biocrust species in isolation and not the community as a whole. Thus, they cannot be scaled to address ecosystem interactions, such as trophic exchanges among biocrust members.

In this study, metatranscriptome sequencing was employed to characterize community gene expression. Biocrusts and below-crust soils were collected from undisturbed sites for comparison to soils that have experienced annual physical trampling for seventeen years. The metatranscriptome sequencing was designed to address three hypotheses. First, that metatranscriptomes can distinguish the activities of surface biocrusts from the below-crust soils. Second, a severe ecosystem disturbance in the form of chronic trampling will significantly alter the metatranscriptome. Finally, a short duration wetting pulse of one hour will be sufficient to capture the initial response of the arid soil communities to a water pulse.

## Material and Methods

### Field site

Samples were collected from the Island of the Sky district of Canyonlands National Park, near Moab, Utah (WGS84, 38°27′27.21″N, 109°32′27.93″W). The study site was located in *Coleogyne ramosissima* (blackbrush) shrubland and harbored biocrusts that differed in development and composition of lichens and cyanobacteria. This site has been employed in a chronic trampling experiment and has been described in detail previously (Kuske et al., 2012; Steven et al., 2015).

The sampling date was October 26^th^ 2012, the most recent previous precipitation event was two days earlier, registering as a trace event (too small to measure) and the year to date precipitation was 6.6 cm, about 35% of the normal year to date average; so the biocrusts were dry at the time of sampling. Samples were collected from the plant interspaces from areas visibly devoid of mosses and lichen to focus on cyanobacteria-dominated biocrusts. The biocrusts were defined as the layer of soil bound together by the cyanobacterial filaments (generally ca. 1 cm depth) and the below-crust soils were the unbound soils directly below at about 2 cm depth.

### Wetting treatment

For wetting, three approximately 13 mm soil diameter cores of ca. 2 cm depth were collected and placed in a Petri dish and 25 ml of sterile distilled water was added from the top, simulating a rainfall. This volume of water is approximately equivalent to a 5 mm precipitation event (based on the volume of the Petri dish). This is within normal precipitation amounts for the region (Hereford and Webb, 1992). The purpose of placing the sample in the Petri dish was to limit the lateral diffusion of water to maintain a consistent wetting treatment between samples. While the Petri dish would not allow for drainage, given the volume of water compared to the volume of biocrust (5 mm water depth compared to 2 cm soil depth) it is unlikely that a significant amount of drainage would have occurred, particularly in the short duration (1 hour) of the wetting pulse.

### Sample collection

Dry samples (*ca.* 10 g) were collected immediately and placed separately into a 50 ml solution of RNA*later* (Sigma-Aldrich) and directly stored on dry ice. For the wetted samples, soil cores were collected and placed in Petri dishes, wetted, and after 1 hour of incubation in the field, the biocrust and below-crust samples were separated and individually placed in RNA*later* and stored on dry ice. After collection, samples were transported frozen to Los Alamos National Laboratory for processing.

### RNA extraction and library preparation

Prior to RNA extraction, each sample was thawed and 4 g of material was removed. RNA was purified using the RNeasy purification kit (Qiagen), and eluted in a final volume of 50 μl. RNA concentration was determined using a Nanodrop spectrophotometer (Thermo Scientific). DNA was removed using Turbo DNase I enzymatic digestion (Life Techonologies). A quantitative real-time PCR (qPCR)-based assay was used to insure that residual DNA was not present in the purified RNA preparations.

Ribosomal RNA (rRNA) was removed from the total RNA via hybridization to biotinylated probes provided in Ribo-Zero kits (EpiCentre). Metabacteria and eukaryote (human/mouse/rat) probes were combined, hybridized to total RNA, and the complexes were removed with the magnetic beads. Samples were processed according to the manufacturer’s protocols.

RNA samples were quantified using a Life Technologies RNA quantification kit and a Qubit 2.0 instrument (Life Technologies). The size distribution of the RNA molecules in each sample was determined using a BioAnalyzer 2100 and RNA pico chips and reagents (Agilent). A maximum of 50 ng of RNA from each sample was converted to an Illumina sequencing library using ScriptSeq kits (EpiCentre). Each library was generated using a unique 6 bp multiplex barcode provided in the ScriptSeq kit. Following library generation, the 12 samples were pooled in equimolar concentrations and run on two lanes of the Illumina HiSeq2000 sequencing platform at the Los Alamos National Laboratory sequencing center, with 2×150 sequence chemistry. Raw sequences were uploaded to the NCBI Sequence Read Archive (SRA) under the Submission ID: SUB3320760 and BioProject ID: PRJNA421954.

### Sequence quality trimming and assembly

Raw sequence reads were quality screened and contaminating sequences were removed with the BBDuk package (v. 37.17) within the BBTools suite (https://jgi.doe.gov/data-and-tools/bbtools/). Briefly, sequence adaptors were removed using the provided adaptors.fa reference file with a minimum kmer match of 11 bp and a 1 bp mismatch allowed for removal. The sequences were then quality trimmed to Q10 using the Phred algorithm with a kmer size of 23 (including the “tbe” and “tbo” flags). Small and large subunit ribosomal RNA sequences that passed the magnetic bead pulldown were bioinformatically removed using BBDuk and the ribokmers.fa reference file (resulting in a removal of 5 to 22% of the sequence reads). Finally, common sequence contaminants were removed with the hg19_main_mask_ribo_animal_allplant_allfungus.fa reference file. Both sequence screenings were performed with a kmer size of 31.

The quality-filtered sequences were assembled with MEGAHIT using default metagenomic parameters (Li et al., 2016). Only contigs greater than 500 base pairs were retained. The number of contigs in the assembly was 209,032 ranging in length from 500 to 35,855 bp The N50 of the assembly was 835 just below the predicted average length of a bacterial mRNA of 924 bp (Xu et al., 2006). These data indicate that many potential full-length gene sequences were recovered. Additionally, contigs >10,000 base pairs suggest that even operon length transcripts were assembled. The assembled contigs are included as a supplemental file to this manuscript (Supplemental File 1).

### Estimation of transcript abundance and taxonomic/functional annotation

The quality filtered raw reads were pseudo-aligned to the assembly to generate transcript abundances in the samples using the kallisto software package (v0.43.1; Bray et al., 2016). First, the assembly was used to generate an index file, using the default parameters (31 kmer length) and the quality trimmed raw read files were mapped to the index file. 100 bootstrap analyses were performed for pseudo replication and import into the Sleuth software package to generate contig count tables (Pimentel et al., 2017).

For sequence annotation contigs were aligned to the NCBI non-redundant (NR) protein database (downloaded July 9^th^, 2017) using DIAMOND with default parameters (Buchfink et al., 2014). Alignments were uploaded to the community version of MEGAN 6 (Huson et al., 2016) and contigs were assigned to SEED subsystems (Overbeek et al., 2014) within MEGAN using the long-read enabled sequence assignment and the May-2015 accession to SEED mapping file. The LCA classification of reads was performed with the prot.accession2taxid mapping file. Merging the contig taxonomic and hierarchical functional annotations with the estimated counts generated a table with LCA classifications and hierarchical SEED annotations for each contig in the assembly and was the raw data on which statistical comparisons were made.

### Statistical analyses

Non-metric multidimensional scaling (NMDS) analyses were performed with the phyloseq R package (McMurdie and Holmes, 2013). Inter-sample distances were calculated with the Bray-Curtis dissimilarity metric, and statistical differences in composition were determined using the PERMANOVA statistic with the adonis function in the vegan R package (Dixon and Palmer, 2003).

To identify significant differences in annotation bins, the annotated count tables were uploaded to the STAMP software package (Parks et al., 2014) along with a relevant metadata table that encoded the information about each sample. For comparisons between multiple groups, differentially abundant bins were identified with an ANOVA test to determine if there was a significant difference in means between the groups, with a p-value cutoff of ≤0.05 and an effect size filter of 0.5. All p-values were corrected for multiple testing using the Bonferroni method. For two group comparisons (i.e. between control/trampled and surface/subsurface) significant differences were identified by Welsh’s t-test with Storey’s FDR method for multiple test correction.

To identify the specific contigs that were differentially abundant due to wetting, unnormalized estimated count data was analyzed with the phyloseq implementation of DESeq2 (Love et al., 2014). Initially, contigs with a summed total count of less than 10 across the samples were removed to avoid the analysis of extremely rare contigs. Differentially abundant contigs were identified with the Wald test on data with a parametric fit. Contigs were considered significantly different in estimated counts if the Benjamin Hochberg multiple test corrected p-values were ≤0.05 with a false detection rate of 0.1. Only contigs showing at least a two-fold change in mean estimated counts were considered biologically meaningful.

## Results

### Clustering of sequence datasets

Differences between the biocrusts and trampled soils were readily apparent at a visual scale (Fig. 1A). To investigate the relationships between the metatranscriptomes from the different samples, the transcript abundances were employed to generate a NMDS plot (Fig. 1B). The control and trampled samples clustered distinctly (PERMANOVA p=0.001), indicating that chronic trampling significantly altered transcript composition (Fig. 1B, large ellipses). The wetting treatment also induced an alteration in transcript composition, as evidenced by the clustering of the wet samples from the dry samples (PERMANOVA p=0.001 for biocrusts, p=0.004 for trampled soils), indicating that gene expression was significantly altered after only one hour of a water pulse (Fig. 1B, inset ellipses).

### Taxonomic bins in metatranscriptomes

Each contig in the assembly was assigned to a taxonomic rank using a lowest common ancestor (LCA) approach, and the estimated counts were used to identify the most common taxa in the datasets. As the LCA method identifies the common taxonomic rank that is shared by all homology matches, sequences are often assigned to broad taxonomic ranks. For example, among the most abundant taxonomic bins in the contigs, were groups such as Cyanobacteria, Bacteria, and cellular organisms (Fig. 2).

Three taxonomic bins accounted for a substantial proportion of the estimated counts, namely the Oscillatoriales an order encompassing many filamentous Cyanobacteria, the phylum Cyanobacteria, and the kingdom Bacteria (Fig. 2). The next most abundant bin was *Microcoleus vaginatus* the specific cyanobacterium that is considered to be a keystone species of biocrusts. Thus, these data suggest that the bulk of RNA in biocrust soils is derived from Cyanobacteria, including the category Bacteria, which could also include sequences derived from Cyanobacteria.

### Functional annotations in the metatranscriptomes

Contigs were functionally annotated to the SEED hierarchy, and the estimated counts were used to investigate the abundance of different functions in the datasets. At the broadest annotations levels the categories Protein Metabolism and Carbohydrates were dominant (Fig. 3). However, for the biocrusts the order was revered with Carbohydrates-related reads being more abundant than for Protein Metabolism (Fig. 3).

To further investigate the most abundant metabolic pathways in the metatranscriptomes, the sequences in the categories Protein Metabolism and Carbohydrates were annotated to the second level in the SEED hierarchy (Fig. 4). Within the category Protein Metabolism the most abundant subsystems were largely involved in structural components of the ribosome or protein synthesis. However, the subsystems also included ATP dependent proteolysis and bacterial proteasomes, indicating a capacity for protein recycling. Only a single subsystem (ribosomal protein S12p) was found to have a significant difference in abundance among the samples.

The most abundant subsystems in the category Carbohydrates were related to carbon fixation (Fig. 5; Calvin-Benson cycle, and alpha/beta carboxysomes) supporting that these soil communities primarily rely on autotrophic carbon fixation. Two central metabolic pathways, the Entner-Doudoroff pathway and TCA cycle were also represented among the most abundant subsystems. Within this category were also subsystems that point to the metabolism of specific carbon compounds, namely maltose, glycerol, and glycogen (Fig. 5). Among the Carbohydrate subsystems, three were identified as significantly different among the metatranscriptomes. Beta-carboxysomes were most abundant in the trampled subsurface soils. Beta-type carboxysomes tend to be more common in heterotrophic bacteria (Kerfeld and Melnicki, 2016), suggesting potential taxonomic shifts in the organisms performing carbon fixation in the trampled subsurface soils. In comparison, Maltose utilization and Photorespiration were both elevated in the wet biocrusts, suggesting increased expression under wetting. Yet, the most abundant subsystems in the datasets tend to be present in similar proportions across the datasets.

### Differentially abundant subsystems due to trampling

Chronic physical trampling induced a significant shift in composition of the metatranscriptomes as evidenced by the NMDS analysis (Fig. 1B). To identify functions underlying the differences, contig annotations were binned to the second level in the SEED hierarchy and differentially abundant functional bins were identified (Fig. 6). A total of 15 functional bins were identified as significantly different. The largest proportional difference was in Nitrogen fixation, which was significantly reduced in the trampled soils (Fig 6). Concurrently, sequence reads to deal with nitrosative stress, the ability to detoxify reactive nitrogen species, was also reduced in the trampled soils, supporting a shift of nitrogen metabolism. Other subsystems point to alterations in amino acid metabolism, specifically methionine and lysine. Two subsystems were also related to the transport and maintenance of metal levels in cells, with a decrease in genes for copper homeostasis as well as nickel and cobalt transport (Fig. 6). Thus, these observations support that chronic trampling alters the metabolism and transport of nitrogen and metals by the soil community.

### Differentially abundant subsystems due to soil depth

The datasets were separated into surface and subsurface samples, irrespective of trampling, to investigate if specific subsystems differentiated the metatranscriptomes based on soil horizon. A total of five significantly different subsystems were identified between surface and subsurface soils (Fig. 7). Of the five subsystems, three were related to the biosynthesis or metabolism of amino acids, and one for the biosynthesis of a carbohydrate polymer, namely arabinogalactan, which consists of arabinose and galactose monosaccharides that can complex with peptidoglycan in the envelope of some bacteria (Crick et al., 2001). Finally, the subsystem for transport of the metals nickel and cobalt was significantly reduced in the subsurface samples, matching an observation for the trampled soils (Fig. 6). Overall, a relatively small number of subsystems discriminated the metatranscriptomes of surface versus subsurface soil communities.

### Differentially abundant contigs due to wetting

Each contig in the assembly was plotted based on its mean abundance and log-fold change between dry and wet samples (Fig. 8). These data clearly demonstrate that the number of contigs that were significantly altered due to wetting was larger in the biocrusts than for any of the other sample types (Fig. 8A, red points). In comparison, no contigs were identified as significantly enriched in the wet trampled surface soils, whereas 27 were significantly enriched in the dry conditions (Fig. 8B). The number of differentially abundant contigs was more synonymous between the below-crust and trampled below soils (Fig. 8C versus 8D).

To investigate the functional role of the differentially abundant contigs, they were annotated to the second level of the SEED hierarchy and the results for the 20 most abundant bins in the biocrusts are summarized in Figure 9. The functional assignments of the contigs split the annotations into three groups: 1. Functions specifically enriched in dry crusts, which included 5-FCL-like proteins (a protein family implicated in thiamin metabolism (Pribat et al., 2011)) and hydrogenases (Fig. 9 top two yellow bars). 2. Functions specifically enriched in wet biocrusts, Photosystem I and II (Fig. 9, bottom two green bars). 3. Functions that included contigs enriched in both the dry and wet biocrusts, which encompassed the remaining functional bins (Fig. 9; Table S1). Results for the surface trampled soils are not displayed as no contigs were significantly enriched in the wetted samples, and of the 27 significantly enriched in the dry samples, only 4 were functionally annotated, all to the category hydrogenases (Table S2).

The below-crust and trampled below-crust samples showed a similar number and pattern of differentially abundant contigs (Fig. 8C,D). Similarities included Hydrogenases, Protein chaperones, Pathogenicity islands, GroEL/GroES, Fe-S-cluster assembly, ATP dependent proteolysis, and Biofilm formation all of which were enriched in the dry soils (Fig. 10). In sum, these observations suggest that while there were substantial differences in the response of the surface soils to the wetting pulse, within the subsurface the effects of wetting were more congruent.

### Biogeochemical cycling genes in the metatranscriptomes

The contig annotations were examined for genes in carbon and nitrogen cycling, of which a reference set were selected to investigate the effect of wetting on biogeochemical cycling (Table 1). As can be seen from Table 1, multiple contigs were mapped to the same annotation, suggesting multiple sequence variants exist of each of the genes. The multiple transcripts for the same gene product is easily explained by multiple organisms encoding the same genes, multiple gene copies, or incomplete assembly of full-length transcripts.

**Table 1.**
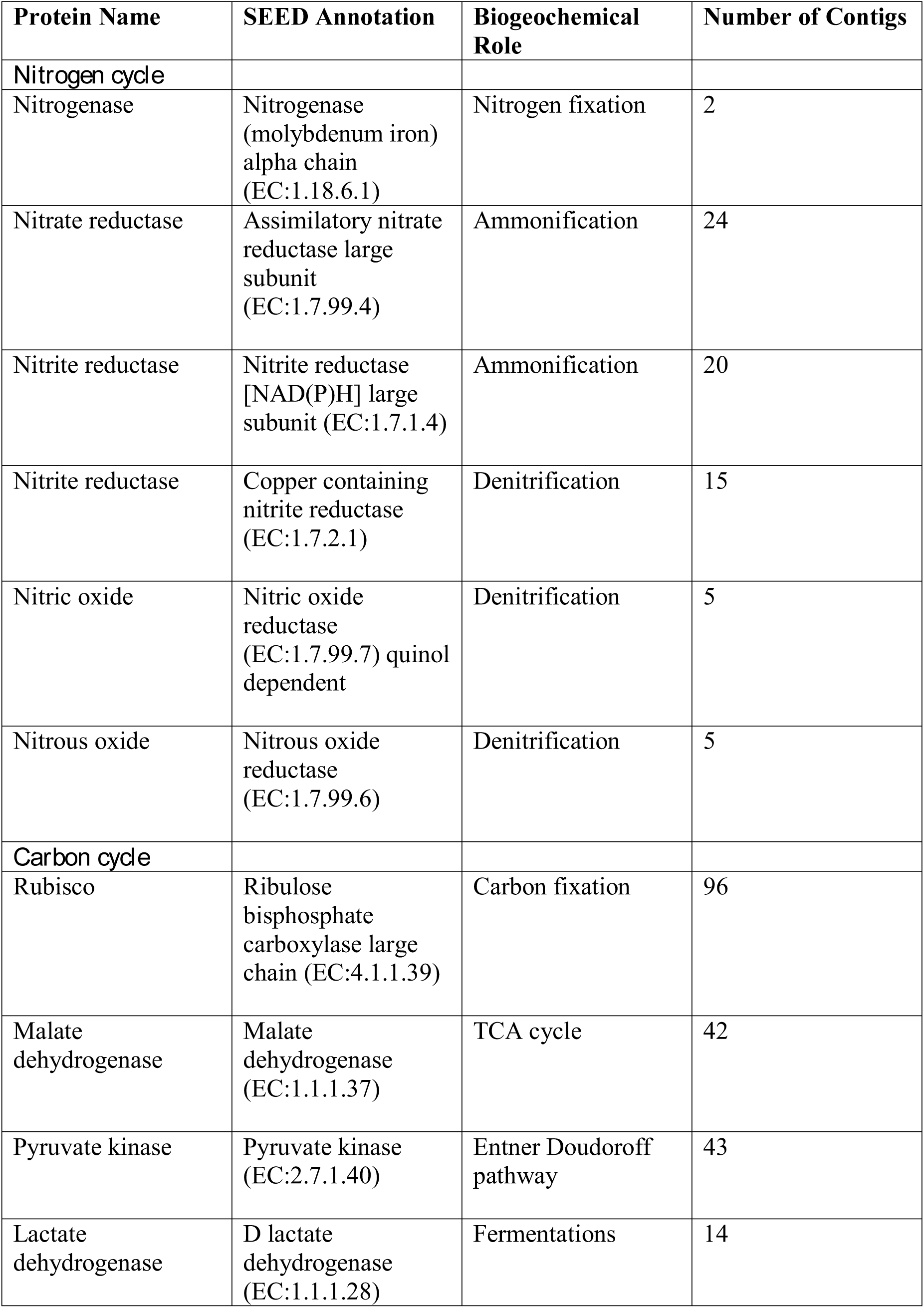
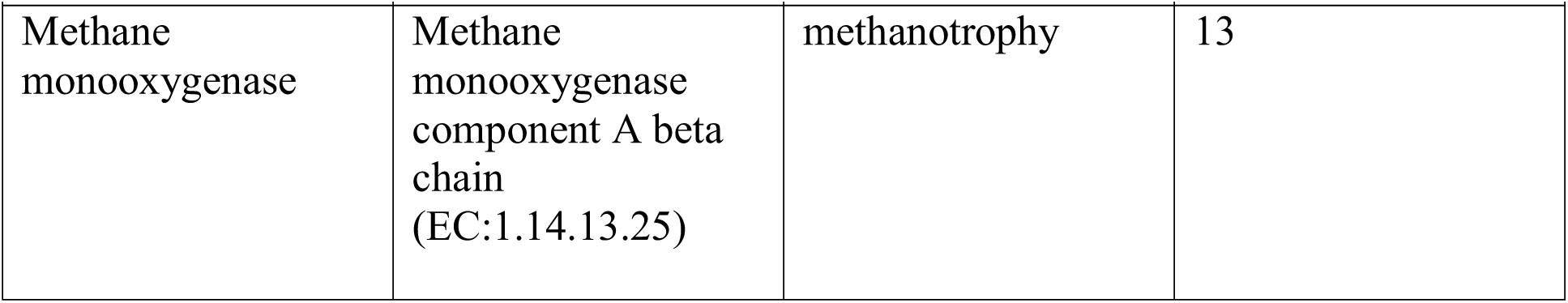
Biogeochemical cycling genes assessed in biocrusts

| **Protein Name** | **SEED Annotation** | **Biogeochemical Role** | **Number of Contigs** |
| --- | --- | --- | --- |
| ***Nitrogen cycle*** |  |  |  |
| Nitrogenase | Nitrogenase (molybdenum iron) alpha chain (EC:1.18.6.1) | Nitrogen fixation | 2 |
| Nitrate reductase | Assimilatory nitrate reductase large subunit (EC:1.7.99.4) | Ammonification | 24 |
| Nitrite reductase | Nitrite reductase [NAD(P)H] large subunit (EC:1.7.1.4) | Ammonification | 20 |
| Nitrite reductase | Copper containing nitrite reductase (EC:1.7.2.1) | Denitrification | 15 |
| Nitric oxide | Nitric oxide reductase (EC:1.7.99.7) quinol dependent | Denitrification | 5 |
| Nitrous oxide | Nitrous oxide reductase (EC:1.7.99.6) | Denitrification | 5 |
| ***Carbon cycle*** |  |  |  |
| Rubisco | Ribulose biphosphate carboxylase large chain (EC:4.1.1.39) | Carbon fixation | 96 |
| Malate dehydrogenase | Malate dehydrogenase (EC:1.1.1.37) | TCA cycle | 42 |
| Pyruvate kinase | Pyruvate kinase (EC:2.7.1.40) | Entner Doudoroff pathway | 43 |
| Lactate dehydrogenase | D lactate dehydrogenase (EC:1.1.1.28) | Fermentations | 14 |
| Methane<br>monooxygenase | Methane<br>monooxygenase<br>component A beta<br>chain<br>(EC:1.14.13.25) | methanotrophy | 13 |

The biogeochemical cycling genes were assessed for their abundance in the dry and wet biocrusts. The biocrusts were chosen as they showed the most pronounced response to the wetting pulse (Fig. 8). Nitrogen cycling genes in the biocrusts did not show a consensus response to wetting. The marker genes for each of the nitrogen cycling processes included contigs that were more abundant in wet and dry conditions, with the exception of nitrogen fixation (Fig. 11). Similarly, carbon cycling genes did not move in concert in response to wetting. In this regard, these data highlight the complexity of the defining a common response of the biocrust community to wetting and linking the changes in transcript abundance to the potential for biogeochemical cycling.

## Discussion

### Metabolic activity in arid soils as revealed by metatranscriptome sequencing

The most abundant functional bins across the metagenomes indicate that central pathways in protein and carbohydrate metabolism account for a relatively large proportion of the metatranscriptomes across the samples (Fig. 3). Broadly similar categories have been described by metagenomic sequencing of biocrust soils, suggesting a correspondence between the most abundant genes and transcripts (Steven et al., 2014, 2015).

The pathways represented among the subsystem Carbohydrates point to specific carbon metabolic processes that support these arid soil communities (Fig. 5). Pathways for the fixation of CO_2_ into biomass are dominant, highlighting the scarcity of organic carbon in these soils and the dependence on autotrophic carbon fixation (Klemmedson, 1989). Beyond carbon fixation, pathways for glycerol, glycogen and maltose metabolism were all abundant in the metatranscriptomes (Fig. 5). Cyanobacteria accumulate intracellular stores of glycogen as a carbon and energy source (De et al., 2017) and maltose can be employed by mixotrophic filamentous cyanobacteria as a heterotrophic carbon source (White and Shilo, 1975). A common strategy employed by microorganisms to survive desiccation is the intracellular accumulation of compatible solutes. These solutes replace water in the cell and aid in maintaining cellular and macromolecule structure (Esbelin et al., 2018). Compatible solutes can take the form of small sugars, glycocides, or peptides (da Costa et al., 1998). A common compatible solute is the sugar trehalose (Hershkovitz et al., 1991; Page-Sharp et al., 1999). *M. vaginatus* encodes genes to synthesize trehalose from both glycogen and maltose (Starkenburg et al., 2011). Thus, glycogen and maltose play a central role in protecting biocrust community members from starvation and desiccation. Thus, the metatranscriptiomes point to a carbon-processing pathway that may be important in supporting the function of biocrusts.

### Metatranscriptomic characterization of an ecosystem disturbance

Biocrust soils subjected to chronic physical trampling were visually distinguishable from healthy biocrusts by the loss of mature developed biocrusts (Fig. 1A). The metatranscriptomes of the trampled soils clustered distinctly by NMDS (Fig. 1B), demonstrated several differentially abundant subsytems (Fig. 6), and displayed a notably altered response to a wetting pulse (Fig. 8). Taken together, these data clearly indicate that chronic physical trampling alters the structure of the biocrust soils and community gene expression.

Trampling of the biocrusts compromises the integrity of the biocrust structure, but some cyanobacteria survive the treatment and can be detected in the soils following the trampling (Kuske et al., 2012; Steven et al., 2015). In this respect, the trampled soils may be comparable to early successional stage biocrusts. In the initial stages of biocrust formation, the founder species *M. vaginatus* is dominant, forming the physical structure of the biocrust. Accordingly, sequences within the bin Oscillatoriales, the order to which *M. vaginatus* belongs, tended to be highest in the trampled datasets (Fig. 2). The dominance of filamentous cyanobacteria in early successional stage crusts comes at a cost. These cyanobacteria are not capable of nitrogen fixation. A lack of diazotrophic cyanobacteria in early and disturbed crusts is supported by reductions of nitrogen fixation potential as measured by acetylene reduction by up to 250% (Evans R. D. and Belnap J., 1999; Zhou et al., 2016). However, the presence of non-cyanobacterial diazotrophs, predominantly within the phyla Firmicutes and Proteobacteria in early successional stage crusts has been presented as evidence that the nitrogen fixation potential of early successional biocrust may be under-estimated (Pepe-Ranney et al., 2016) The metatranscriptome data presented here suggest that nitrogen fixation potential is significantly reduced in trampled soils as assessed by the abundance of nitrogen fixation transcripts in the trampled soils (Fig. 6). However, these data only consider dry soils and a brief wetting pulse; perhaps the wetting threshold for activity of diazotrophic organisms in trampled soils is larger and thus requires a longer or larger wetting pulse. Yet, these data could also go some way to explain why biocrusts were generally unresponsive to inorganic nitrogen additions (Mueller et al., 2015). Perhaps a more robust response would be observed with nitrogen addition to disturbed or early-successional stage biocrusts, where nitrogen fixation potential may be limited.

In addition to reduced nitrogen fixation, the trampled surface soils also displayed a dramatic difference in the number of contigs that were altered in abundance due to wetting in comparison to the biocrusts (Fig. 8A versus 8B). No contigs were significantly enriched in the wet trampled surface soils in comparison to more than 4000 for the biocrusts. Biocrusts produce an exopolysaccharide matrix and dark pigments that play a strong role in retaining moisture (Mazor et al., 1996; Rossi et al., 2012). Well-developed biocrusts also alter the soil surface properties, increasing roughness and forming pinnacles, properties that can additionally foster moisture retention and infiltration (Belnap, 2006). These properties are visually absent in the trampled soils (Fig. 1A), presumably also affecting the ability of the community to take advantage of a wetting pulse. In this respect, these data suggest that not only are the trampled soil communities altered in their metabolic repertoire, but are less able to take advantage of the pulsed wetting events that control the productivity of these arid soils.

### Differences between surface and subsurface metatranscriptomes

One of the unexpected findings of this study was the relatively small difference in the metatranscriptomes between the surface and subsurface samples. Because of gradients in resources such as light and moisture, biocrust communities are vertically stratified, often at the millimeter scale (Garcia-Pichel et al., 2003; Steven et al., 2013). Yet, only a small number of functional bins were identified as significantly different between the surface and subsurface samples (Fig. 7). The most parsimonious explanation for these results is that the cyanobacteria in both soil layers accounted for most of the transcriptome. The below-crust and trampled subsurface soils both showed a dominance of sequence belonging to cyanobacteria (Fig. 2), although metagenomic sequencing and other metrics indicate heterotrophic bacteria and Achaea are dominant in the subsurface soils (Steven et al., 2013). The high proportion of cyanobacteria in these soils can be easily explained by an incomplete separation of the biocrusts and the below-crust soils, or more likely that the small proportion of cyanobacteria that do inhabit the sub-crust soils account for most of the transcriptional activity. These data highlight the need to look at longer term pulsed experiments to investigate if and when other members of the below-crust soils become active.

### Response of arid soil communities to a wetting pulse

In general, mRNA molecules are considered to be short-lived and of modest abundance in the cell. For example, physiological studies of *Escherichia coli* suggest that actively growing cells contain approximately 1,000 mRNA molecules with a half-life of about five minutes (Bernstein et al., 2002). The mRNA content of bacterial cells in the open ocean is more modest, with approximately 200 mRNA transcripts per cell, also with half-lives in the range of a few minutes (Moran et al., 2013). However, transcriptional activity in active microbial cultures or other ecosystems may not be an appropriate comparison to biocrusts. Arid lands are characterized by extended dry periods, during which biocrust respiration is virtually undetectable suggesting that biocrusts are largely dormant during this time (Garcia-Pichel and Belnap, 1996; Huxman et al., 2004; Sponseller, 2007). Periods of metabolic activity, stimulated primarily by water availability, occur infrequently and are short in duration. In this regard, biocrust response to wetting may be more similar to the germination of plant seeds. Plant seeds are dormant (or quiescent) until the initiation of germination, which is triggered by the uptake of water (Nonogaki et al., 2010). The resumption of metabolic activity for plant seeds (or arid soil microbes) is dependent on mRNA produced by the organism prior to the period of dormancy. In seeds mRNA transcripts are protected as RNA/protein complexes, also referred to as mRNPs (Bewley, 1997; Reichel et al., 2016). A similar mechanism could serve to protect transcripts in dry biocrusts, allowing them to be stable over the duration of dry periods. Several cyanobacteria encode functional homologs to eukaryotic RNA-binding proteins (Mulligan et al., 1994). The specific functions of these genes have not been conclusively demonstrated, but they appear to play a role in cold-stress or nitrogen starvation (Mori et al., 2003; Sugita et al., 1999). Here we propose another potential role for cyanobacteria mRNPs; that is protecting mRNA molecules from desiccation damage during biocrust drying. The observation of complex pools of mRNA transcripts in dry conditions across the soil types, including transcripts more abundant in the dry samples (Figs. 4,5,7), points to mechanisms for maintaining RNA integrity in dry arid soils. The rapid turnover of mRNA is also supported by the differences in the metatranscriptomes after only one hour of wetting, particularly for the biocrusts (Fig. 8). This matches other observations that short-term transcriptional responses may be overlooked in linking transcriptional data to ecosystem process rates (Albright et al.). Future work investigating the lifespan of arid soil mRNA molecules may shed light transcript stability as a regulator of community responses to desiccation and rewetting.

A common observation across the dry samples was an enrichment of transcripts for hydrogenases (Fig. 9,10), specifically uptake hydrogenases (see supplemental tables). Uptake hydrogenases are bi-directional, capable of both producing and consuming H_2._ Uptake hydrogenases have been found among all heterocystous cyanobacteria studied to date, as the enzyme captures H_2_ produced during nitrogen-fixation, although non-nitrogen fixing cyanobacteria also contain uptake hydrogenases (Tamagnini et al., 2005). Several cyanobacteria within the order Oscillatoriales have been found to be prodigious producers of H_2_, although H_2_ evolution was absent from *M. vaginatus* (Kothari et al., 2012). Thus, these transcriptomic data point to cyanobacteria possibly producing and/or consuming H_2_ in dry conditions. However, the majority of the uptake hydrogenase transcripts enriched in the dry soils were found from various Actinobacteria, specifically the Solirubrobacterales (see supplemental tables). Scavenging of atmospheric hydrogen has been proposed as a mechanism by which bacteria can maintain metabolic activity under low carbon availability (Greening et al., 2015). Thus an active H_2_ based metabolism appears to be operating in the dry biocrusts that may involve a give and take between the Cyanobacteria and Actinobacteria.

By the metric of the number of differentially abundant transcripts, the wetting pulse was most pronounced in the biocrust soils (Fig. 8). Many of the subsystems to which the contigs were annotated were within central metabolic pathways and machinery, such a translation elongation factors, ATP synthase, ribosomal proteins, and protein chaperones (Fig. 9).

In the biocrusts most of the differentially abundant subsystems were made up of contigs with members that were identified as both more abundant in the wet and dry conditions. For example, contigs annotated to bacterial small subunit (SSU) ribosomal proteins (Fig. 9). It is well established that bacteria scale their ribosomal counts with metabolic activity (Elser et al., 2003; Steven et al., 2017). In the dry crusts the enriched contigs for SSU ribosomal proteins were primarily from groups of Actinobacteria whereas SSU genes enriched in the wet crusts were primarily cyanobacterial in origin (Table S2). Thus, these data point to different populations being active in the wet and dry conditions. Specifically, this is consistent with cyanobacteria becoming more active in the wet biocrusts. Concurrently, an enrichment of photosystem I, II, and phycobillisome genes supports a rapid recovery of the cyanobacteria and the initiation of photosynthesis (Fig. 9).

### Metatranscriptomes and biocrust biogeochemical cycling

A large driver for studying the microbial ecology of soils is to include soil microorganisms in ecosystem, carbon, and climate models (Reid, 2012). This requires that transcriptional abundances correlate with ecological process rates, a relationship that has not been well established (Moran et al., 2013; Prosser, 2015). In this study we focused on carbon and nitrogen fixation, given the large role biocrusts play in these processes in arid ecosystems (Barger et al., 2016; Sancho et al., 2016). These metatranscriptome datasets highlight two significant challenges in relating transcript abundance to biogeochemical cycling. First, complex communities encode multiple isoforms of the transcripts for proteins in the selected biogeochemical pathways (Table 1; Fig. 11). Multiple transcripts arise from functionally redundant species encoding the same pathways, as well as the potential for a single genome to encode multiple gene copies that may be differentially regulated under different environmental conditions (Allison and Martiny, 2008; Misra and Tuli, 2000). As a result all transcripts in a pathway do not respond in concert.

Another potential confounding factor linking biogeochemical cycling gene transcript abundances to process rates is an observation discussed previously. A significant response to the wetting was that particular bacteria increased their transcripts for ribosomal proteins, presumably leading to elevated cellular ribosomal content (Fig. 9). Because mRNA molecules are not consumed during translation, and in bacteria multiple ribosomes can translate the same mRNA molecule in polysomes, an increased ribosome content can lead to more proteins being produced from the same number of mRNA molecules (Scott et al., 2010). In this manner, cells could increase the stock of biogeochemical cycling proteins through increased ribosomal content without a concurrent increase in the mRNA encoding those genes. In this regard, linking transcript abundance to ecosystem process rates will also need to account for factors such as ribosome content, RNA degradation, and translation efficiency (Kerkhof and Ward, 1993; Pedersen, 1984; Taylor et al., 2013).

## Conclusion

The data presented here suggest that metatranscriptome sequencing can identify differences in biocrust communities in response to perturbations that occur over dramatically different time scales. Chronic physical trampling carried out over 17 years, alters the metabolic potential of the community as well as shaping the response of the community to a wetting pulse. In contrast, a short durational wetting pulse of 1 hour is also sufficient to alter the metatranscriptome. The biocrust communities respond rapidly to a pulse of water, with the cyanobacteria ramping up protein synthesis and photosynthesis genes. Taken together, these observations support that chronic trampling induces shifts in the structure and function of the soil communities that could not be assessed at a visual scale alone, which is often used to asses biocrust health and function (e.g. Belnap et al., 2008; Miller, 2008; Ferrenberg et al., 2015), thus highlighting the importance for the role of molecular microbial ecology tools in characterizing biocrusts and the response and resiliency to ecosystem disturbances.

## Acknowledgements

This study was supported by a U. S. Department of Energy, Biological and Environmental Research (BER) Science Focus Area award to CRK. In addition, BS was partially supported by the Los Alamos Laboratory Directed Research and Development (LDRD) program. Field experiments were initiated and maintained through U.S. DOE Terrestrial Ecosystem Science grants to Jayne Belnap of the U.S. Geological Survey, the National Park Service, and the USGS Climate and Land Use and Ecosystem programs. Any use of trade names is for descriptive purposes only and does not imply endorsement by the U.S. Government. The authors would also like to thank Robert Bjornson from the Yale Center for Research Computing for technical assistance with computational analyses, La Verne Gallegos-Graves for sample preparation at the Los Alamos National Laboratory, and Michaeline Nelson for editorial comments.

## Figure Legends

**Figure 1**. A) Photograph of field plots. The control plots show well-developed biocrusts directly adjacent to a trampled plot (separated by a nylon rope). A pen is included for scale. B). NMDS plot showing the relationship of metatranscriptome datasets. Intergroup distances were calculated with the Bray-Curtis metric. The colored ellipses are included as a visual aid only and were not part of the analysis. The groupings within each ellipse are indicated by the bolded text. The stress value of the ordination is indicated in the top left corner.

**Figure 2.** Taxonomic bins in the metatranscriptomes. Contigs were assigned a lowest common ancestor (LCA) and binned based on these assignments. Estimated counts of the contigs were then employed to assign a percent of total estimated counts for each LCA bin, of which the 20 most abundant are displayed. Each point represents the mean abundance from three replicate samples.

**Figure 3.** SEED functional bins in the metatranscriptomes. Contigs were annotated within the SEED hierarchy and binned based to the broadest categorical assignments (level 1). Estimated counts of the contigs were then employed to assign a percent of total estimated counts for each bin, of which the 20 most abundant are displayed. Each point represents the mean abundance from three replicate samples.

**Figure 4.** Subsystems within the category Protein Metabolism. The 20 most abundant level 2 subsystems within the category are shown. Sample abbreviations: CD=crust dry; CW=crust wet; BCD=below-crust dry; BCW=below-crust wet; TSD=Trampled surface dry; TSW=Trampled surface wet; TBD=Trampled below dry; TBW=Trampled below wet. The size of the points represent the proportion of estimated counts as a percent of the total reads in the parent subsystem (i.e. percent of Protein Metabolism reads). Each point represents the mean of three replicate samples. Points colored red were significantly different by an ANOVA difference of means test.

**Figure 5.** Subsystems within the category Carbohydrates. The 20 most abundant level 2 subsystems within the category are shown. Sample abbreviations: CD=crust dry; CW=crust wet; BCD=below-crust dry; BCW=below-crust wet; TSD=Trampled surface dry; TSW=Trampled surface wet; TBD=Trampled below dry; TBW=Trampled below wet. The size of the points represent the proportion of estimated counts as a percent of the total reads in the parent subsystem (i.e. percent of Carbohydrate reads). Each point represents the mean of three replicate samples. Points colored red were significantly different by an ANOVA difference of means test.

**Figure 6.** Differentially abundant subsystems due to trampling. Boxplot of differentially abundant subsystems between biocrust and trampled soils. For this analysis the wet and dry, as well as the surface and subsurface samples were considered together. Thus, the boxes summarize the data for 12 replicate samples. For each box the mean is represented by the bolded bar and boxes represent the interquartile range. Outliers are shown as points outside of the box and whiskers.

**Figure 7.** Differentially abundant subsystems due to soil depth. Boxplot of differentially abundant subsystems between surface and subsurface soils. For this analysis the wet and dry, as well as the biocrust and trampled samples were considered together. Thus, the boxes summarize the data for 12 samples. For each box the mean is represented by the bolded bar and boxes represent the interquartile range. Outliers are shown as points outside of the box and whiskers.

**Figure 8.** Bland-Altman plots displaying shifts in contig abundance due to wetting. The average contig abundance (log_10_ scale) is displayed on the x-axis. As the means are for both wet and dry samples, each point is the average of six samples. The fold-change (log_2_ scale) is displayed on the y-axis, such that contigs enriched in dry samples are shown below the 0 line and contigs enriched in wet are shown above. Points having a significant difference with False Detection Rates (FDR) of less than 0.1 and at least a two-fold difference in abundance (indicated by the light gray dashed lines) are colored red.

**Figure 9.** Differentially abundant subsystems due to wetting in surface soil samples. The level 2 SEED annotations were used to functionally bin the contigs identified as differentially abundant (Fig. 8), of which the 20 most abundant (sorted by the total number of contigs in each bin) are displayed. A full list of differentially abundant contigs in the biocrusts is listed in supplemental Table S1 (only annotated contigs are presented). For the trampled surface samples, no contigs were identified as significantly enriched in the wetted samples (Fig. 8) and of the 27 significantly enriched in the dry samples only four were annotated to SEED subsystems, so the data is not shown. All four subsystems were in the category Hydrogenases. A full list of differentially abundant contigs in the trampled surface soils is listed in supplemental Table S2.

**Figure 10.** Differentially abundant subsystems due to wetting in subsurface soil samples. The level 2 SEED annotations were used to functionally bin the contigs identified as differentially abundant (Fig. 8). A) Below-crust soils, B) Below trampled soils. A full list of differentially abundant contigs in the below-crust soils is listed in supplemental Table S3 and in supplemental Table S4 for the below trampled soils.

**Figure 11.** Biogeochemical cycling genes in the biocrust metatranscriptomes. The biogeochemical cycling genes listed in Table 1 were mapped onto the biocrust datasets. Each point represents the average normalized abundance of each contig in the dataset (n=6). Biogeochemical cycling genes are colored based on the biogeochemical pathway to which they belong.

## References

Abed, R. M. M., Polerecky, L., Al-Habsi, A., Oetjen, J., Strous, M., and de Beer, D. (2014). Rapid Recovery of Cyanobacterial Pigments in Desiccated Biological Soil Crusts following Addition of Water. PLoS ONE 9, e112372. doi:10.1371/journal.pone.0112372.

Albright, M. B.., Johansen, R., Lopez, D., Gallegos-Graves, L., Steven, B., Kuske, C. R., et al. Short-term transcriptional response of microbial communities to N-fertilization in pine forest soil. Appl. Environ. Microbiol.

Allison, S. D., and Martiny, J. B. (2008). Resistance, resilience, and redundancy in microbial communities. Proc. Natl. Acad. Sci. 105, 11512–11519.

Barger, N. N., Weber, B., Garcia-Pichel, F., Zaady, E., and Belnap, J. (2016). “Patterns and Controls on Nitrogen Cycling of Biological Soil Crusts,” in Biological Soil Crusts: An Organizing Principle in Drylands, eds. B. Weber, B. Büdel, and J. Belnap (Cham: Springer International Publishing), p257–285. doi:10.1007/978-3-319-30214-0_14.

Belnap, J. (2003). The world at your feet: desert biological soil crusts. Front. Ecol. Environ. 1, 181–189.

Belnap, J. (2006). The potential roles of biological soil crusts in dryland hydrologic cycles. Hydrol. Process. 20, 3159–3178. doi:10.1002/hyp.6325.

Belnap, J., Phillips, S. L., Witwicki, D. L., and Miller, M. E. (2008). Visually assessing the level of development and soil surface stability of cyanobacterially dominated biological soil crusts. J. Arid Environ. 72, 1257–1264. doi:10.1016/j.jaridenv.2008.02.019.

Belnap, J., Weber, B., and Büdel, B. (2016). “Biological Soil Crusts as an Organizing Principle in Drylands,” in Biological Soil Crusts: An Organizing Principle in Drylands Ecological Studies. (Springer, Cham), 3–13. doi:10.1007/978-3-319-30214-0_1.

Bernstein, J. A., Khodursky, A. B., Lin, P.-H., Lin-Chao, S., and Cohen, S. N. (2002). Global analysis of mRNA decay and abundance in Escherichia coli at single-gene resolution using two-color fluorescent DNA microarrays. Proc. Natl. Acad. Sci. 99, 9697–9702. doi:10.1073/pnas.112318199.

Bewley, J. D. (1997). Seed germination and dormancy. Plant Cell 9, 1055.

Bray, N. L., Pimentel, H., Melsted, P., and Pachter, L. (2016). Near-optimal probabilistic RNA-seq quantification. Nat. Biotechnol. 34, 525–527. doi:10.1038/nbt.3519.

Buchfink, B., Xie, C., and Huson, D. H. (2014). Fast and sensitive protein alignment using DIAMOND. Nat. Methods 12, 59–60. doi:10.1038/nmeth.3176.

Crick, D. C., Mahapatra, S., and Brennan, P. J. (2001). Biosynthesis of the arabinogalactan-peptidoglycan complex of Mycobacterium tuberculosis. Glycobiology 11, 107R–118R.

da Costa, M. S., Santos, H., and Galinski, E. A. (1998). “An overview of the role and diversity of compatible solutes in Bacteria and Archaea,” in Biotechnology of Extremophiles, ed. G. Antranikian (Berlin, Heidelberg: Springer Berlin Heidelberg), 117–153. doi:10.1007/BFb0102291.

De, A. P., Frigaard, N.-U., and Sakuragi, Y. (2017). Determination of the Glycogen Content in Cyanobacteria. J. Vis. Exp. JoVE.

Dixon, P., and Palmer, M. W. (2003). VEGAN, a package of R functions for community ecology. J. Veg. Sci. 14, 927–930. doi:10.1658/1100-9233(2003)014[0927:VAPORF]2.0.CO;2.

Elser, J. J., Acharya, K., Kyle, M., Cotner, J., Makino, W., Markow, T., et al. (2003). Growth rate-stoichiometry couplings in diverse biota. Ecol. Lett. 6, 936–943. doi:10.1046/j.1461-0248.2003.00518.x.

Esbelin, J., Santos, T., and Hébraud, M. (2018). Desiccation: An environmental and food industry stress that bacteria commonly face. Food Microbiol. 69, 82–88. doi:10.1016/j.fm.2017.07.017.

Evans R. D., and Belnap J. (1999). Long-term consequences of disturbance on nitrogen dynamics in an arid ecosystem. Ecology 80, 150–160. doi:10.1890/0012-9658(1999)080[0150:LTCODO]2.0.CO;2.

Evans, R. D., and Lange, O. L. (2001). “Biological soil crusts and ecosystem nitrogen and carbon dynamics,” in Biological Soil Crusts: Structure, Function, and Management Ecological Studies., eds. J. Belnap and O. L. Lange (Springer Berlin Heidelberg), p263–279. Available at: http://link.springer.com/chapter/10.1007/978-3-642-56475-8_20 [Accessed February 19, 2015].

Ferrenberg, S., Reed, S. C., and Belnap, J. (2015). Climate change and physical disturbance cause similar community shifts in biological soil crusts. Proc. Natl. Acad. Sci. 112, 12116–12121. doi:10.1073/pnas.1509150112.

Gao, B., Zhang, D., Li, X., Yang, H., and Wood, A. J. (2014). De novo assembly and characterization of the transcriptome in the desiccation-tolerant moss Syntrichia caninervis. BMC Res. Notes 7, 490.

Garcia-Pichel, F., and Belnap, J. (1996). Microenvironments and Microscale Productivity of Cyanobacterial Desert Crusts1. J. Phycol. 32, 774–782. doi:10.1111/j.0022-3646.1996.00774.x.

Garcia-Pichel, F., Johnson, S. L., Youngkin, D., and Belnap, J. (2003). Small-Scale Vertical Distribution of Bacterial Biomass and Diversity in Biological Soil Crusts from Arid Lands in the Colorado Plateau. Microb. Ecol. 46, 312–321. doi:10.1007/s00248-003-1004-0.

Garcia-Pichel, F., and Pringault, O. (2001). Microbiology: Cyanobacteria track water in desert soils. Nature 413, 380–381. doi:10.1038/35096640.

Greening, C., Carere, C. R., Rushton-Green, R., Harold, L. K., Hards, K., Taylor, M. C., et al. (2015). Persistence of the dominant soil phylum Acidobacteria by trace gas scavenging. Proc. Natl. Acad. Sci. 112, 10497–10502. doi:10.1073/pnas.1508385112.

Hereford, R., and Webb, R. H. (1992). Historic variation of warm-season rainfall, Southern Colorado Plateau, Southwestern U.S.A. Clim. Change 22, 239–256. doi:10.1007/BF00143030.

Hershkovitz, N., Oren, A., and Cohen, Y. (1991). Accumulation of Trehalose and Sucrose in Cyanobacteria Exposed to Matric Water Stress. Appl. Environ. Microbiol. 57, 645–648.

Huson, D. H., Beier, S., Flade, I., Górska, A., El-Hadidi, M., Mitra, S., et al. (2016). MEGAN Community Edition-Interactive Exploration and Analysis of Large-Scale Microbiome Sequencing Data. PLOS Comput. Biol. 12, e1004957. doi:10.1371/journal.pcbi.1004957.

Huxman, T. E., Snyder, K. A., Tissue, D., Leffler, A. J., Ogle, K., Pockman, W. T., et al. (2004). Precipitation pulses and carbon fluxes in semiarid and arid ecosystems. Oecologia 141, 254–268. doi:10.1007/s00442-004-1682-4.

Karaoz, U., Couradeau, E., da Rocha, U. N., Lim, H.-C., Northen, T., Garcia-Pichel, F., et al. (2018). Large Blooms of Bacillales (Firmicutes) Underlie the Response to Wetting of Cyanobacterial Biocrusts at Various Stages of Maturity. mBio 9, e01366–16.

Kerfeld, C. A., and Melnicki, M. R. (2016). Assembly, function and evolution of cyanobacterial carboxysomes. Curr. Opin. Plant Biol. 31, 66–75. doi:10.1016/j.pbi.2016.03.009.

Kerkhof, L., and Ward, B. B. (1993). Comparison of Nucleic Acid Hybridization and Fluorometry for Measurement of the Relationship between RNA/DNA Ratio and Growth Rate in a Marine Bacterium. Appl. Environ. Microbiol. 59, 1303–1309.

Klemmedson, J. O. (1989). Soil organic matter in arid and semiarid ecosystems: Sources, accumulation, and distribution. Arid Soil Res. Rehabil. 3, 99–114. doi:10.1080/15324988909381194.

Kothari, A., Potrafka, R., and Garcia-Pichel, F. (2012). Diversity in hydrogen evolution from bidirectional hydrogenases in cyanobacteria from terrestrial, freshwater and marine intertidal environments. J. Biotechnol. 162, 105–114. doi:10.1016/j.jbiotec.2012.04.017.

Kuske, C. R., Yeager, C. M., Johnson, S., Ticknor, L. O., and Belnap, J. (2012). Response and resilience of soil biocrust bacterial communities to chronic physical disturbance in arid shrublands. ISME J. 6, 886–897. doi:10.1038/ismej.2011.153.

Li, D., Luo, R., Liu, C.-M., Leung, C.-M., Ting, H.-F., Sadakane, K., et al. (2016). MEGAHIT v1.0: A fast and scalable metagenome assembler driven by advanced methodologies and community practices. Methods 102, 3–11. doi:10.1016/j.ymeth.2016.02.020.

Love, M. I., Huber, W., and Anders, S. (2014). Moderated estimation of fold change and dispersion for RNA-seq data with DESeq2. Genome Biol. 15, 550. doi:10.1186/s13059-014-0550-8.

Mazor, G., Kidron, G. J., Vonshak, A., and Abeliovich, A. (1996). The role of cyanobacterial exopolysaccharides in structuring desert microbial crusts. FEMS Microbiol. Ecol. 21, 121–130.

McMurdie, P. J., and Holmes, S. (2013). phyloseq: an R package for reproducible interactive analysis and graphics of microbiome census data. PloS One 8, e61217.

Miller, M. E. (2008). Broad-Scale Assessment of Rangeland Health, Grand Staircase–Escalante National Monument, USA. Rangel. Ecol. Manag. 61, 249–262. doi:10.2111/07-107.1.

Misra, H. S., and Tuli, R. (2000). Differential Expression of Photosynthesis and Nitrogen Fixation Genes in the Cyanobacterium Plectonema boryanum. Plant Physiol. 122, 731–736.

Moran, M. A., Satinsky, B., Gifford, S. M., Luo, H., Rivers, A., Chan, L.-K., et al. (2013). Sizing up metatranscriptomics. ISME J. 7, 237–243. doi:10.1038/ismej.2012.94.

Mori, S., Castoreno, A., Mulligan, M. E., and Lammers, P. J. (2003). Nitrogen status modulates the expression of RNA-binding proteins in cyanobacteria. FEMS Microbiol. Lett. 227, 203–210. doi:10.1016/S0378-1097(03)00682-7.

Mueller, R. C., Belnap, J., and Kuske, C. R. (2015). Soil bacterial and fungal community responses to nitrogen addition across soil depth and microhabitat in an arid shrubland. Front. Microbiol. 6. doi:10.3389/fmicb.2015.00891.

Mulligan, M. E., Jackman, D. M., and Murphy, S. T. (1994). Heterocyst-forming Filamentous Cyanobacteria Encode Proteins that Resemble Eukaryotic RNA-binding Proteins of the RNP Family. J. Mol. Biol. 235, 1162–1170. doi:10.1006/jmbi.1994.1070.

Nonogaki, H., Bassel, G. W., and Bewley, J. D. (2010). Germination—Still a mystery. Plant Sci. 179, 574–581. doi:10.1016/j.plantsci.2010.02.010.

Overbeek, R., Olson, R., Pusch, G. D., Olsen, G. J., Davis, J. J., Disz, T., et al. (2014). The SEED and the Rapid Annotation of microbial genomes using Subsystems Technology (RAST). Nucleic Acids Res. 42, D206–D214. doi:10.1093/nar/gkt1226.

Page-Sharp, M., Behm, C. A., and Smith, G. D. (1999). Involvement of the compatible solutes trehalose and sucrose in the response to salt stress of a cyanobacterial Scytonema species isolated from desert soils. Biochim. Biophys. Acta BBA - Gen. Subj. 1472, 519–528. doi:10.1016/S0304-4165(99)00155-5.

Parks, D. H., Tyson, G. W., Hugenholtz, P., and Beiko, R. G. (2014). STAMP: statistical analysis of taxonomic and functional profiles. Bioinformatics 30, 3123–3124. doi:10.1093/bioinformatics/btu494.

Pedersen, S. (1984). Escherichia coli ribosomes translate in vivo with variable rate. EMBO J. 3, 2895–2898.

Pepe-Ranney, C., Koechli, C., Potrafka, R., Andam, C., Eggleston, E., Garcia-Pichel, F., et al. (2016). Non-cyanobacterial diazotrophs mediate dinitrogen fixation in biological soil crusts during early crust formation. ISME J. 10, 287–298. doi:10.1038/ismej.2015.106.

Pimentel, H., Bray, N. L., Puente, S., Melsted, P., and Pachter, L. (2017). Differential analysis of RNA-seq incorporating quantification uncertainty. Nat. Methods 14, 687.

Pointing, S. B., and Belnap, J. (2012). Microbial colonization and controls in dryland systems. Nat. Rev. Microbiol. 10, 551–562. doi:10.1038/nrmicro2831.

Pribat, A., Blaby, I. K., Lara-Núñez, A., Jeanguenin, L., Fouquet, R., Frelin, O., et al. (2011). A 5-formyltetrahydrofolate cycloligase paralog from all domains of life: comparative genomic and experimental evidence for a cryptic role in thiamin metabolism. Funct. Integr. Genomics 11, 467–478. doi:10.1007/s10142-011-0224-5.

Pringault, O., and Garcia-Pichel, F. (2004). Hydrotaxis of Cyanobacteria in Desert Crusts. Microb. Ecol. 47. doi:10.1007/s00248-002-0107-3.

Prosser, J. I. (2015). Dispersing misconceptions and identifying opportunities for the use of “omics” in soil microbial ecology. Nat. Rev. Microbiol. 13, 439. doi:10.1038/nrmicro3468.

Rajeev, L., Da Rocha, U. N., Klitgord, N., Luning, E. G., Fortney, J., Axen, S. D., et al. (2013). Dynamic cyanobacterial response to hydration and dehydration in a desert biological soil crust. ISME J. 7, 2178–2191.

Reichel, M., Liao, Y., Rettel, M., Ragan, C., Evers, M., Alleaume, A.-M., et al. (2016). In planta determination of the mRNA-binding proteome of Arabidopsis etiolated seedlings. Plant Cell, tpc.p00562.2016. doi:10.1105/tpc.16.00562.

Reid, A. (2012). Incorporating Microbial Processes into Climate Models. Available at: http://www.asmscience.org/content/report/colloquia/colloquia.19 [Accessed November 16, 2017].

Rossi, F., Potrafka, R. M., Pichel, F. G., and De Philippis, R. (2012). The role of the exopolysaccharides in enhancing hydraulic conductivity of biological soil crusts. Soil Biol. Biochem. 46, 33–40. doi:10.1016/j.soilbio.2011.10.016.

Sancho, L. G., Belnap, J., Colesie, C., Raggio, J., and Weber, B. (2016). “Carbon Budgets of Biological Soil Crusts at Micro-, Meso-, and Global Scales,” in Biological Soil Crusts: An Organizing Principle in Drylands Ecological Studies. (Springer, Cham), 287–304. doi:10.1007/978-3-319-30214-0_15.

Scott, M., Gunderson, C. W., Mateescu, E. M., Zhang, Z., and Hwa, T. (2010). Interdependence of Cell Growth and Gene Expression: Origins and Consequences. Science 330, 1099–1102. doi:10.1126/science.1192588.

Sponseller, R. A. (2007). Precipitation pulses and soil CO 2 flux in a Sonoran Desert ecosystem. Glob. Change Biol. 13, 426–436. doi:10.1111/j.1365-2486.2006.01307.x.

Starkenburg, S. R., Reitenga, K. G., Freitas, T., Johnson, S., Chain, P. S. G., Garcia-Pichel, F., et al. (2011). Genome of the Cyanobacterium Microcoleus vaginatusFGP-2, a Photosynthetic Ecosystem Engineer of Arid Land Soil Biocrusts Worldwide. J. Bacteriol. 193, 4569–4570. doi:10.1128/JB.05138-11.

Steven, B., Gallegos-Graves, L. V., Belnap, J., and Kuske, C. R. (2013). Dryland soil microbial communities display spatial biogeographic patterns associated with soil depth and soil parent material. FEMS Microbiol. Ecol. 86, 101–113. doi:10.1111/1574-6941.12143.

Steven, B., Gallegos-Graves, L. V., Yeager, C. M., Belnap, J., and Kuske, C. R. (2014). Common and distinguishing features of the bacterial and fungal communities in biological soil crusts and shrub root zone soils. Soil Biol. Biochem. 69, 302–312.

Steven, B., Hesse, C., Soghigian, J., Gallegos-Graves, L. V., and Dunbar, J. (2017). Simulated rRNA/DNA Ratios Show Potential To Misclassify Active Populations as Dormant. Appl. Environ. Microbiol. 83, e00696–17. doi:10.1128/AEM.00696-17.

Steven, B., Kuske, C. R., Reed, S. C., and Belnap, J. (2015). Climate Change and Physical Disturbance Manipulations Result in Distinct Biological Soil Crust Communities. Appl. Environ. Microbiol., AEM. 01443–15.

Sugita, C., Mutsuda, M., Sugiura, M., and Sugita, M. (1999). Targeted deletion of genes for eukaryotic RNA-binding proteins, Rbp1 and Rbp2, in the cyanobacterium Synechococcus sp. strain PCC7942: Rbp1 is indispensable for cell growth at low temperatures. FEMS Microbiol. Lett. 176, 155–161. doi:10.1111/j.1574-6968.1999.tb13656.x.

Tamagnini, P., Leitão, E., and Oxelfelt, F. (2005). Uptake hydrogenase in cyanobacteria: novel input from non-heterocystous strains. Biochem. Soc. Trans. 33, 67–69. doi:10.1042/BST0330067.

Taylor, R. C., Webb Robertson, B.-J. M., Markille, L. M., Serres, M. H., Linggi, B. E., Aldrich, J. T., et al. (2013). Changes in Translational Efficiency is a Dominant Regulatory Mechanism in the Environmental Response of Bacteria. Integr. Biol. Quant. Biosci. Nano Macro 5, 1393–1406. doi:10.1039/c3ib40120k.

Tucker, C. L., McHugh, T. A., Howell, A., Gill, R., Weber, B., Belnap, J., et al. (2017). The concurrent use of novel soil surface microclimate measurements to evaluate CO2 pulses in biocrusted interspaces in a cool desert ecosystem. Biogeochemistry 135, 239–249. doi:10.1007/s10533-017-0372-3.

White, A. W., and Shilo, M. (1975). Heterotrophic growth of the filamentous blue-green alga Plectonema boryanum. Arch. Microbiol. 102, 123–127.

Xu, L., Chen, H., Hu, X., Zhang, R., Zhang, Z., and Luo, Z. W. (2006). Average Gene Length Is Highly Conserved in Prokaryotes and Eukaryotes and Diverges Only Between the Two Kingdoms. Mol. Biol. Evol. 23, 1107–1108. doi:10.1093/molbev/msk019.

Zhou, X., Smith, H., Silva, A. G., Belnap, J., and Garcia-Pichel, F. (2016). Differential Responses of Dinitrogen Fixation, Diazotrophic Cyanobacteria and Ammonia Oxidation Reveal a Potential Warming-Induced Imbalance of the N-Cycle in Biological Soil Crusts. PLOS ONE 11, e0164932. doi:10.1371/journal.pone.0164932.

